# Extreme temperature exposure has negative demographic consequences for *Sulfolobus acidocaldarius*

**DOI:** 10.64898/2026.09.26.754634

**Authors:** CD Durrant, Rebecca Lowry Palmer, Danna R Gifford

**Affiliations:** Division of Evolution, Infection and Genomics, School of Biological Sciences, Faculty of Biology, Medicine and Health, The University of Manchester

**Keywords:** thermophilic archaea, heat stress, thermal tolerance, extreme temperature, environmental fluctuations, geothermal springs, stress recovery, *Sulfolobus acidocaldarius*

## Abstract

Microorganisms inhabiting geothermal springs and volcanic systems experience fluctuating temperatures that can periodically exceed their upper thermal limits, but the demographic consequences of such exposure remain poorly understood. Here, we investigated demographic responses of the thermophilic archaeon *Sulfolobus acidocaldarius* to an extreme temperature (94.1°C) under two regimes: sustained exposure varying in duration, and episodic exposure interspersed with recovery at a permissive temperature (75 °C). Under sustained exposure, populations showed no detectable loss of viability after 15 min but declined thereafter, decreasing by approximately five orders of magnitude after 120 min. Under episodic exposure, populations remained viable across nine exposure-recovery cycles but declined in density with successive cycles. Similar responses were observed for three strains, including a DNA mismatch repair knockout (Δ*nucS*), indicating that mismatch repair deficiency did not affect viability or recovery. Together, these results demonstrate that *S. acidocaldarius* can withstand brief and repeated exposure to near-boiling temperatures, with mortality determined primarily by cumulative exposure duration rather than a fixed thermal threshold.

**Importance:** This study shows that microbial tolerance to extreme temperature depends not only on the temperature reached, but also on how long and how often that stress occurs. In *Sulfolobus acidocaldarius*, brief near-boiling exposure was tolerated, whereas prolonged or repeated exposure reduced population recovery. This contrasts with previous experimental work in which repeated heat exposure promoted adaptation, showing that recurrent thermal stress does not necessarily lead to increased tolerance. These findings highlight the importance of temporal environmental variation in shaping survival and show why fixed thermal limits alone may not capture microbial persistence in fluctuating extreme environments.

## Introduction

Microorganisms inhabiting geothermal springs experience pronounced spatial and temporal variation in temperature. Convection, fluid mixing and pulsed hydrothermal discharge continually shift thermal conditions, exposing microorganisms to extreme temperatures for different durations and at different frequencies (1,2). Cells may therefore encounter temperatures above their growth range either as isolated events, or as repeated exposures separated by periods permitting growth. Whether populations persist will depend not only on the temperature reached, but also on how long exposure lasts and whether cells can recover between events. Yet the demographic consequences of these contrasting exposure regimes remain poorly understood.

The thermoacidophilic archaeon *Sulfolobus acidocaldarius* is a well-established model for life at high temperature, with a reported growth range of approximately 65 –85 °C (3). Yet the geothermal systems it inhabits can contain temperatures approaching boiling, with gradients from permissive to lethal conditions changing over spatial scales of only a few metres (4,5). While it has been established that *S. acidocaldarius* can survive briefly at these extreme temperatures through physiological changes (3,6), how its survival and subsequent population recovery is affected by the duration and recurrence of extreme-temperature exposures is unexplored. We therefore examined how exposure duration and exposure recurrence affect survival and recovery at a near-boiling temperature (*T*_extreme_ = 94.1°C). Populations underwent either one continuous exposure of varied duration, or multiple ‘episodic’ short exposures separated by recovery periods. These treatments approximate two forms of thermal variation in geothermal springs: sustained near-boiling conditions near hydrothermal upwellings or active vents (5,7), and recurrent high-temperature events, for example, those associated with geyser eruption–recharge cycles (8).

In microbes, loss of DNA mismatch repair (MMR) can produce mutator phenotypes that may accelerate adaptation by generating beneficial mutations, but may also reduce fitness through deleterious mutation accumulation (9–13). Archaea generally lack the canonical MMR homologs of bacteria and eukaryotes, instead possessing the endonuclease NucS (also called EndoMS), which functions analogously (14). Inactivation of NucS produces mutator phenotypes in *Saccharolobus islandicus* and actinobacteria (15–20). *S. acidocaldarius* NucS corrects the same mismatches *in vitro* (21), but the *in vivo* role is unresolved: no mutator phenotype was observed using a locus reporter assay (21), but one biased towards indels (22), which NucS is now known not to correct (17). Consequently, the genome-wide effect of NucS in *S. acidocaldarius* may be underappreciated. We therefore investigated whether inactivation of NucS, a putative functional analog of canonical MMR, altered survival during or recovery after extreme-temperature exposure.

A single brief exposure to *T*_extreme_ caused no detectable loss of viability, but longer exposures progressively reduced viability by several orders of magnitude and reduced post-recovery population density after a fixed recovery period. Episodic short exposures also progressively decreased post-recovery population density with increasing numbers of episodes. NucS inactivation had no detectable effect under either exposure regime, providing no evidence that this putative MMR system influenced extreme temperature survival or recovery over the timescale examined. Together, these results indicate that the persistence of *S. acidocaldarius* at extreme temperature events comes at a demographic cost and depends on the opportunity for growth recovery between events.

## Methods

### Strains

Three strains of *S. acidocaldarius* were used, DSM639, SK1, and SK7. DSM639 is a wild-type strain first isolated in 1972 at Locomotive Spring within the Norris Geyser Basin of Yellowstone National Park, USA (originally designated isolate 98-3 by Brock *et al*. in 1972 (23)). SK1 possesses a knockout of restriction endonuclease gene *suaI*, resulting in loss of restriction activity against unmethylated DNA (24). SK7 is a derivative of SK1 and carries a further deletion of DNA mismatch-repair endonuclease gene *nucS* (21). Strains SK1 and SK7 are derived from a spontaneous uracil auxotroph, MR31 (*pyrE-13*, (25)); MR31 is ultimately derived from strain DG1, which is genomically similar to DSM639 (26).

### Growth conditions

Strains were cultured in Brock’s basal media (BBM+: 1 mL/L of Brock I, 10 mL/L of Brock II/III, and 1 mL/L of 0.1 M FeCl_3_·6H_2_O, (23)) supplemented with 10 mL/L each of filter-sterilised 30% D-glucose, 10% N-Z-amine and 0.18 mM filter-sterilised uracil (SK1 and SK7 are uracil auxotrophs). Final pH was adjusted to 3 with ∼150 µl/L of H_2_SO_4_.

Culturing was performed as described in Al-Baqasami *et al*. (2024) (27). Briefly, cultures were grown in 1.5 mL BBM+ in 2 mL microcentrifuge tubes with pierced lids. Gas-permeable membranes (Breathe-Easy, Sigma-Aldrich, Germany) were placed over pierced lids to allow oxygen diffusion. Incubation was performed in thermomixers (DLAB Thermo Mix, DLAB, China) with 300 rpm shaking. Transfers into fresh medium were performed by inoculating 15 μL of culture into pre-heated 1.5 mL BBM+, corresponding to a 10^−2^ dilution. Negative controls of uninoculated BBM+ were used in all experiments. Cultured in this way, populations typically reach stationary phase within 48 h, with an initial density of approximately 10^7^ cells and a final density of 10^9^ cells, with a linear relationship between cell counts and blank-corrected OD_600_ observed across the range 0.03 –1.0 (27).

### Sustained extreme temperature varying in duration

To determine how viability and recovery change with the duration of extreme heat exposure, cultures of *S. acidocaldarius* were grown for 4 d at 75 °C and subsequently exposed to extreme temperature for varying lengths of time. Thermomixers were preheated to a set point of 95 °C, although the culture medium reached a maximum measured temperature of 94.1 °C, hereafter defined as *T*_extreme_. Exposure time was measured from the transfer of cultures from 75 °C to the preheated thermomixers and therefore included the heating period. Cultures took an average of 4 min 49 s to reach 90.3 °C and 6 min 22 s to reach 94.1 °C. Temperatures above 90 °C are known to reduce viability and induce the heat shock response in *S. acidocaldarius* (3,6,28). All durations reported include heating time.

To assess viability, cultures of DSM639 were exposed to *T*_extreme_ for 15 –120 min. Following exposure, cultures were serially diluted ten-fold and spot plated onto solid medium. To assess recovery after exposure, cultures of DSM639, SK1 and SK7 were exposed to *T*_extreme_ for 15 –330 min. Following exposure, cultures were diluted 10^−2^ into fresh BBM+ and incubated at 75 °C for a further four days. Population density was assessed using blank-corrected OD_600_. This experiment was repeated twice with two populations of each strain per time point.

### Episodic short-duration extreme temperature

To investigate population responses to episodic extreme heat, cultures were subjected to repeated exposures to *T*_extreme_ = 94.1°C, separated by recovery periods at the optimal growth temperature. Cultures of *S. acidocaldarius* were grown for 4 d at 75 °C and then heated and exposed to *T*_extreme_ for 15 min. Following each exposure, cultures were diluted 10^-2^ into fresh BBM+ medium and returned to 75 °C for a subsequent 4 d. Previous work has shown that populations can withstand recurrent 10^−2^ dilution and regrowth at 75 °C without declining (27). This exposure–recovery cycle was performed nine times using serially transferred populations. Population density (blank-corrected OD_600_) was measured immediately before each exposure, following the preceding growth or recovery period. The 15-min exposure duration was selected because a single exposure did not noticeably reduce viability. The experiment comprised three independent repeats, each containing seven replicate populations per strain.

### Data analysis

All statistical analyses were performed in R (version 4.3.1) (29). For the sustained duration experiment, effects of time and strain on log10(OD_600_) were assessed using linear modelling with time as a continuous covariate and strain as a discrete factor. For the episodic experiment, effects of episode and strain on log10(OD_600_) were assessed using linear mixed-effects modelling using the lmer function from lme4 (version 1.1-33) and lmerTest (version 3.1-3) packages (30,31). Episode was treated as a continuous covariate, strain as a discrete fixed effect and replicate population as a random effect. Post hoc contrasts on estimated marginal means used Dunnett’s *p*-value adjustment for multiple testing.

## Results and Discussion

Our experiments provide insight into how *S. acidocaldarius* maintains population viability in thermally variable geothermal environments. Changes in hydrothermal discharge and mixing with cooler groundwater or surface water create steep thermal gradients that shift through space and time, potentially exposing populations to brief temperatures above their growth optimum (3,23,32).

### Viability and recovery declines with increased exposure time to extreme temperature

To determine how the duration of extreme-temperature exposure affected survival of *S. acidocaldarius*, we measured both colony-forming ability immediately after exposure and demographic recovery at a permissive temperature. No apparent reduction in viability was observed after 15 min, but longer exposures progressively reduced survival, producing approximate 100-fold, 10,000-fold and 100,000-fold reductions after an additional 30, 60 and 120 min, respectively (all times inclusive of 6 min 22 s heating time to reach *T*_extreme_ = 94.1°C, **Figure 1A**). Population density after regrowth at 75 °C also declined with increasing exposure duration (linear model, slope = −0.33±0.07 S.E., *t*_127_ = −4.56, *p* = 1.2×10^−5^, **Figure 1B**). There was no significant effect of strain or interaction between strain and time(p > 0.05). This indicates a demographic cost of prolonged extreme-temperature events, with longer exposures producing lower population densities after a fixed-length recovery period. Moreover, recovery was rarely observed following exposures longer than 180 min, suggesting a limit to the duration of near-boiling exposure from which populations can resume growth under these conditions.

**Figure 1:**
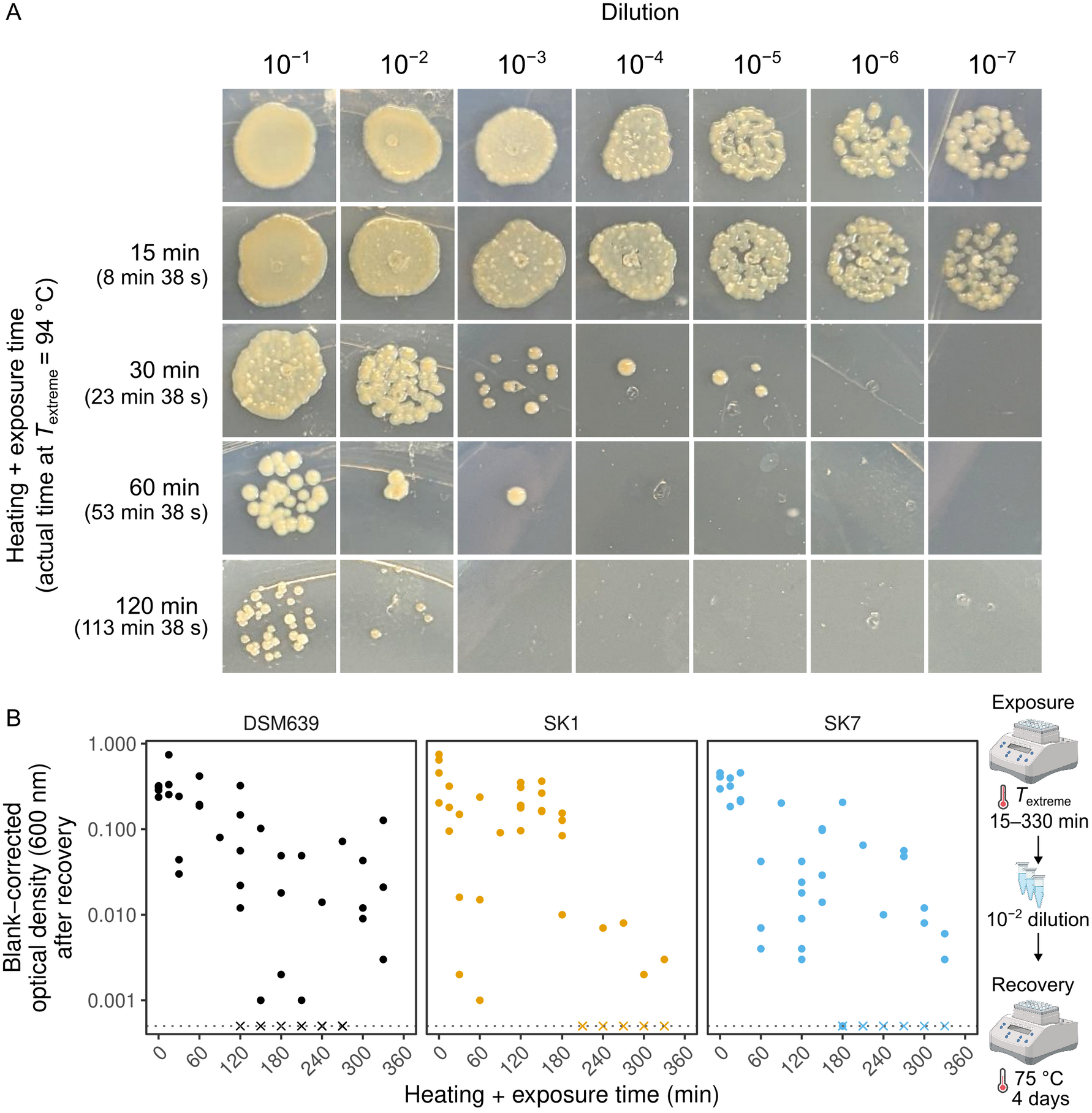
Viability and recovery of S. acidocaldarius populations exposed to extreme temperature for varying durations. A) Viability of stationary phase DSM639 following heating and exposure to *T*_extreme_ (94.1 °C) for the times indicated. B) Recovery of population density (OD_600_) following heating and exposure to *T*_extreme_ for 15 –330 min and subsequent regrowth at 75 °C for four days. Points represent replicate populations for each strain (DSM639, SK1, SK7; *n* = 3 replicates for each duration); crosses indicate measurements below the limit of detection (dotted horizontal line).

The progressive reduction in viability and recovery with increased exposure time suggests a broad distribution of times to death within the population, rather than an abrupt loss of viability at a common exposure duration. Two non-exclusive processes could underlie this distribution: stochastic accumulation of lethal damage and physiological or heritable variation in heat resistance. Both have been implicated in bacterial thermal inactivation (33–36).

Archaeal studies also support thermotolerance as a plastic trait: preconditioning increased *Saccharolobus shibatae* survival at 95 °C, gradual warming extended the thermal growth range of *Metallosphaera sedula*, and heating rate affected *S. acidocaldarius* survival (3,37,38). Notably, our cultures remained viable longer than in Baes *et al*. (2020) (3), who observed total loss of viability after 30 min at 93.5 °C. Baes *et al*. used exponentially growing cultures, in contrast to ours, which were at stationary phase. The difference may simply reflect greater initial density in our cultures, which had approximately ten-fold colony forming units.

However, a more interesting possibility is that nutrient limitation in stationary phase could increase thermotolerance through membrane remodelling. Both starvation and elevated temperatures favour increased GDGT cyclisation, which is crucial for high-temperature growth (39). Whether nutrient limitation enhances extreme-temperature survival warrants further investigation, as such cross-protection could benefit *S. acidocaldarius* in the nutrient-poor geothermal environments it inhabits.

### Episodic short-term exposure to extreme temperature is associated with gradual demographic decline

To assess the consequences of episodic short-duration extreme-temperature exposure, we measured population density across nine exposure–recovery episodes (**Figure 2**). Although a single brief exposure caused no detectable reduction in immediate viability (**Figure 1**), post-recovery density declined with episode number (linear mixed-effects model: slope = −0.11±0.03 S.E., *t*_177_ = −3.22, *p* = 0.00153). Neither strain nor the interaction between strain and episode number had a significant effect (*p* > 0.05).

**Figure 2:**
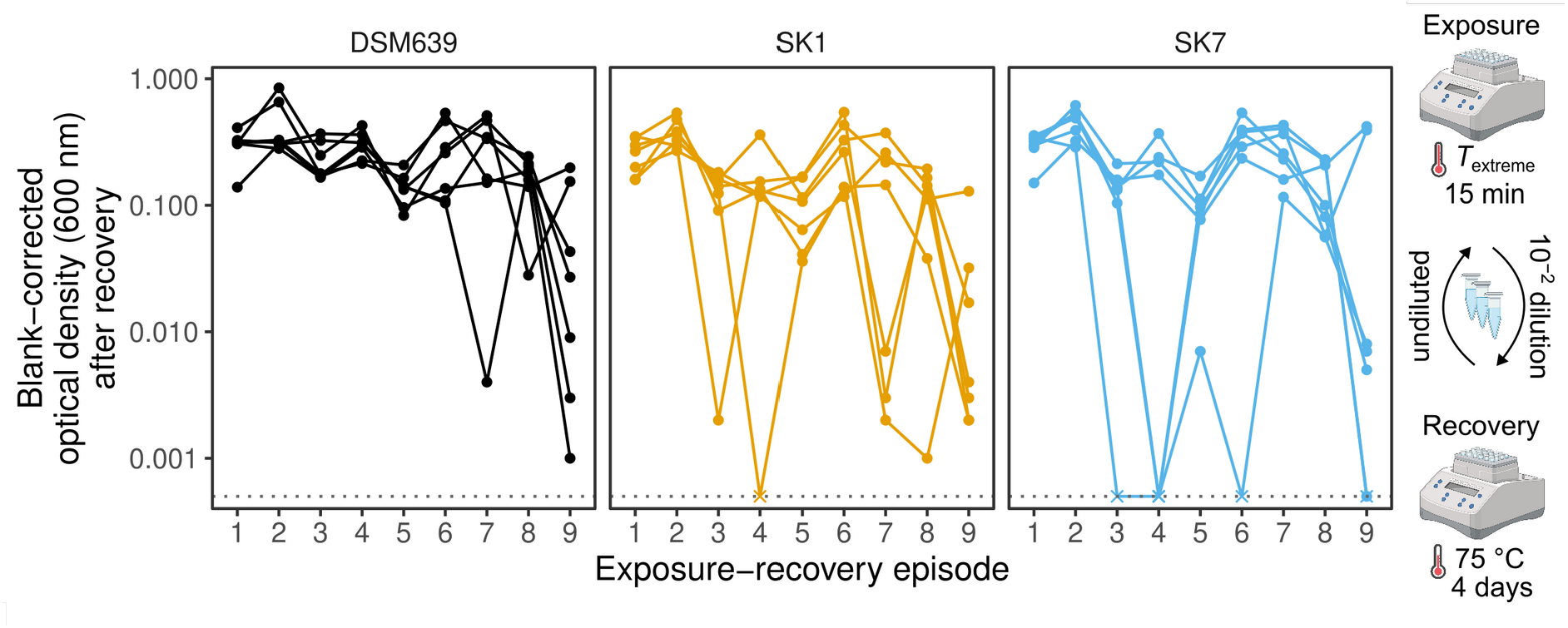
Population density (OD_600_) of S. acidocaldarius populations experiencing episodic short-term extreme temperature exposure followed by recovery. Populations were exposed to *T*_extreme_ (94.1°C) for 15 min, followed by regrowth at 75 °C for four days. Points connected by lines represent repeated measures on replicate populations for each strain (DSM639, SK1, SK7; *n* = 7 of each strain); crosses indicate measurements below the limit of detection (dotted horizontal line).

These results show that tolerance of an isolated short exposure does not necessarily make recurrent exposures demographically neutral. The progressive decline is unlikely to reflect damage accumulating within the same cells over multiple episodes, as the 100-fold dilution and regrowth following each episode means that the population mostly comprises cells produced during the intervening recovery periods. Instead, the decline could arise from population-level carry-over, whereby incomplete recovery left fewer cells to seed each subsequent cycle, or from stress-induced physiological states inherited by descendants. Alternatively, history-dependent phenotypes could in principle persist across generations through inherited proteins, protein aggregates, metabolic states or regulatory feedback, as observed in several studies (40–42).

The progressive decline in post-recovery density contrasts with the response observed in other exposure–recovery experimental evolution studies. Mortier et al. (2021) used a strikingly similar exposure–recovery design: stationary-phase cultures underwent nine 15-min heat shocks at a lethal temperature, each followed by 10^-2^ dilution and regrowth (43). However, in contrast to our results, they observed an increase in survival probability over time, and no decrease in permissive growth. Mall et al. (2026) performed a similar experiment in *Methylobacterium extorquens* with 20 episodes, where adaptation shortened lag time during the recovery phase (44). The contrasting outcomes suggest that demographic decline is not an inevitable consequence of episodic extreme-temperature exposure. Why adaptation was not apparent for our populations remains unresolved, but several possibilities are worth exploring. Adaptation may not have occurred at all, due to limited opportunity for beneficial mutations to arise and spread or because few genetic changes can improve survival at a temperature approaching the upper limit of physiological tolerance. Alternatively, adaptation may have affected components not revealed by post-recovery density—for instance, if antagonistic pleiotropy between survival at *T*_extreme_reduces growth under permissive temperatures.

### No detectable effect of inactivating NucS-mediated DNA mismatch repair

In microbes, loss of DNA MMR can impair acute-stress responses through deleterious mutation load or promote adaptation during recurrent exposure by increasing mutation supply (9–14), but we found no evidence for that effect here. There was no significant difference in the effect of exposure time on recovery from acute exposure [difference in slopes (SK7−SK1) = 0.073±0.10 S.E., *t*_123_ = 0.70, *p* = 0.70]. Thus, we found no evidence that NucS inactivation produced a deleterious mutational load sufficient to compromise recovery from different exposure times. Similarly, recovery across successive extreme-temperature episodes did not differ between the strains [difference in slopes (SK7−SK1) = 0.047±0.05 S.E., *t*_177_ = 0.99, *p* = 0.51], providing no evidence that NucS inactivation affected demographic recovery during recurrent thermal stress.

The absence of a NucS-inactivation effect could be that its role in DNA repair is limited in vivo, as suggested by Suzuki and Kurosawa (2019) (21). Alternatively, even if NucS inactivation substantially increases the mutation rate, adaptation may remain limited because the resulting mutations are either insufficient in number or poorly matched to the genetic changes required to overcome the demographic effects of extreme-temperature exposure. Our work does not exclude a role for mutation rate more generally; NucS is only one of several mutation avoidance systems in *S. acidocaldarius* (45).

### Implications for the dispersal and persistence of *S. acidocaldarius* in natural environments

Together, these experiments show how the temporal structure of thermal variation may influence the ecological distribution of *S. acidocaldarius*. Brief exposure to near-boiling temperature permitted survival and subsequent regrowth, whereas prolonged or recurrent exposure imposed substantial demographic costs. Hotter microhabitats may therefore act as filters on dispersal: cells transported through them may remain viable when transit is brief, but longer residence could reduce successful migration and impose population bottlenecks. Such bottlenecks could reduce within-population diversity while increasing drift-driven differentiation among populations. Anderson et al. (2017) similarly invoked bottlenecks, albeit through a different proximal mechanism, to explain the lower genomic diversity in wild isolates of *S. acidocaldarius* (46). Temporal temperature variation provides a complementary mechanism through which environmental instability may shape population persistence and diversification. Notably, natural populations occur at considerably lower densities than those examined here (47–49), so equivalent thermal mortality may leave fewer survivors and reinforce the effects of genetic drift.

## Data availability statement

Data are deposited in a public repository (https://doi.org/10.48420/33838966), which was made available at the point of peer review.

## Acknowledgements

The authors thank Professor Norio Kurosawa (Soka University) for strains SK1 and SK7, and Dr Zahraa Al-Baqasami (Kuwait University) for technical advice. This work was supported by grants to DRG from UKRI-NERC (NE/X012662/1) and The Royal Society (RGS\R1\231308).

## Author contributions

**CD:** Methodology; Investigation; Formal analysis; Data curation; Visualization; Writing – original draft; Writing – review & editing.

**RLP:** Investigation; Methodology; Supervision; Visualization; Writing – review & editing.

**DRG:** Conceptualization; Methodology; Formal analysis; Data curation; Visualization; Supervision; Project administration; Funding acquisition; Writing – review & editing.

